# A Knock-in Ntsr1-Flp Driver Enables Intersectional and Systemic Targeting of Heterogeneous Midbrain Dopamine Circuits

**DOI:** 10.64898/2026.03.11.711093

**Authors:** Fernando Garcia, Andrew P. Villa, Justine T. Wong, Renqi Wu, Priscilla Coriano, Lief E. Fenno, Gina M. Leinninger, Andrew D. Steele

## Abstract

Precise genetic access to molecularly defined neuronal subpopulations is essential for dissecting circuit heterogeneity. We developed and validated a knock-in *neurotensin receptor 1* (*Ntsr1*)*^Flp^*mouse line for intersectional targeting of Ntsr1-expressing neurons. Following delivery of Flp-dependent adeno-associated viral (AAV) reporters, robust recombination was observed throughout the the midbrain and several other brain regions. A subset of Ntsr1^+^ neurons in the substantia nigra and ventral tegmental area lacked dopaminergic markers, indicating that midbrain Ntsr1 populations comprise both dopaminergic and non-dopaminergic neurons. Systemic delivery of a Cre and Flp-dependent reporter in complementary dual-recombinase configurations revealed configuration-dependent differences in dopaminergic targeting. Cis-gene controls (*Dat^Cre^*;*Dat^Flp^*) defined the maximal dopaminergic targeting ceiling and helped distinguish true non-dopaminergic targeting from recombinase-dependent off-target labeling. Finally, a dual-recombinase-dependent taCaspase-3 construct enabled selective ablation of midbrain dopamine neurons in vivo, establishing *Ntsr1^Flp^* as a versatile driver for scalable Boolean targeting.

## Introduction

The peptide Neurotensin (Nts) acts via the G-protein coupled receptors neurotensin receptor-1 (Ntsr1) and neurotensin receptor-2 (Ntsr2). Intriguingly, administration of Nts into different brain regions causes wildly varying effects, from modulating feeding, drinking, movement, social behavior, thermoregulation, cardiovascular control and analgesia.^1–4^ This is likely because Ntsr-expressing target cells vary in neurochemistry and connectivity, such that Nts signaling via them produces very distinct effects. Hence, the challenge of studying the Nts system is characterizing how the brain site- and molecularly-distinct subsets of Ntsr-expressing cells modify physiology and behavior. It is now recognized that Ntsr2 is primarily expressed on astrocytes and endothelial cells, which are being investigated for Nts-Ntsr2 mediated roles in analgesia and cerebrovascular regulation.^5^ By contrast, Ntsr1 is expressed by specific subsets of neurons throughout the brain.^6^ This has spurred the field to characterize site-specific Ntsr1-expressing populations and match their specific roles within the array of Nts-mediated physiology.

One site of interest is the midbrain, where Ntsr1 has emerged as a critical marker of physiological relevance.^6^ Midbrain dopamine neurons are the primary arbiters of motivation,^7^ reinforcement learning,^8,9^ movement,^10,11^ reward,^12,13^ reward-prediction error,^14^ and are also required for drinking and feeding.^15,16^ Far from being a monolithic population, these neurons comprise transcriptionally and functionally heterogeneous subtypes that are often defined by their unique expression of GPCRs and neuropeptides. Single-cell profiling has underscored this diversity, revealing that specific molecular signatures dictate how dopaminergic neuron sub-populations process distinct environmental and internal cues.^17–19^ Among these, the Neurotensin receptor 1 (Ntsr1) has emerged as a critical marker of physiological relevance.^20^ As a high-affinity receptor for the tridecapeptide neurotensin, and a potent modulator of reward and satiety, Ntsr1 identifies a large subset of neurons that sit at the intersection of energy balance and dopaminergic signaling.^21–23^ Understanding how these molecularly defined subsets of dopaminergic neurons coordinate complex behaviors requires genetic tools capable of isolating Ntsr1-expressing cells from the broader midbrain milieu.

Within the midbrain, Ntsr1 marks two regionally defined populations: a substantial subset of VTA (also called A10) dopaminergic neurons implicated in feeding regulation and diet-induced obesity, and Ntsr1 is also expressed in most SN (also called A9) dopaminergic neurons.^1,23–26^ While the VTA and SN both contain dopaminergic neurons, the VTA is typically thought of in modulating motivated behavior while the SN canonically modulates movement, although it has also recently been implicated in motivational or habitual behaviors. Neurotensin signaling modulates dopaminergic neuron excitability and reward-related behaviors, yet it remains unclear whether the behavioral functions attributed to Ntsr1 arise specifically from dopaminergic neurons or from mixed neuronal populations, and whether they differ between the VTA and SN. Although an *Ntsr1^Cre^* line is available,^21^ Cre-based access alone limits experimental flexibility when intersectional targeting with other widely used Cre driver lines is desired. Dual recombinase strategies using Cre and Flp enable Boolean intersectional targeting of molecularly defined neuronal populations^27^ and because many Cre driver lines already exist, complementary Flp knock-in lines substantially expand the range of possible genetic intersections. Prior to this work, however, no Flp-based Ntsr1 knockin line was available, limiting flexible intersectional access to Ntsr1-defined circuits and scalable systemic AAV approaches that require a high fidelity Flp driver to faithfully recapitulate endogenous expression patterns. Furthermore, such a line is well suited for use with newly engineered systemic AAVs,^28,29^ enabling brain-wide intersectional access without invasive intracranial injections.

The increasing recognition of transcriptional and functional heterogeneity within midbrain dopaminergic systems has exposed limitations of single-driver genetic strategies. Cre-only approaches, while powerful, often capture molecularly overlapping but functionally distinct neuronal populations, complicating causal interpretation of behavioral manipulations. Dual-recombinase strategies enable Boolean intersectional logic, permitting selective access to neurons defined by combinatorial gene expression rather than a single marker.^27,30^ In the context of emerging systemic AAV technologies and enhancer-driven viral tools, the availability of high-fidelity Flp driver lines is essential for scalable and precise targeting of molecularly defined neuronal subsets. Thus, generation of an *Ntsr1^Flp^* allele fills a critical methodological gap, enabling refined dissection of Ntsr1-defined circuits within heterogeneous populations.

Here, we describe the development and validation of a knock-in *Ntsr1^Flp^*recombinase mouse line. We demonstrate robust Flp-dependent recombination following stereotaxic delivery of Flp-dependent (fDIO) AAV reporters and show compatibility with *dopamine transporter* (*Slc6a3*, *Dat*)*^Cre^*in dual-recombinase configurations to control fluorescent labeling and genetically encoded cell ablation. Additionally, we introduce a novel Cre- and Flp-dependent taCaspase-3-TEVp construct to enable precise intersectional cell ablation.^31,32^ Together, these experiments establish the *Ntsr1^Flp^* line as a versatile platform for both local and systemic intersectional targeting of dopaminergic and non-dopaminergic Ntsr1 populations in the midbrain.

## Results

### Creation of P2A linked Ntsr1-Flp allele and physiological characterization of Ntsr1^FlpO^ mice

To enable intersectional targeting of Ntsr1-expressing neurons, we generated an Ntsr1^Flp^ knock-in mouse line by inserting a 2A-FlpO cassette into the endogenous *Ntsr1* locus (ENSMUSG00000027568) (Figure 1A). The cassette was positioned in-frame within the Ntsr1 coding sequence to preserve endogenous gene regulation while enabling bicistronic expression of FlpO. CRISPR/Cas9-mediated targeting at exon 4 was confirmed by 5′ and 3′ junctional PCR in G1 offspring, with founder G0 serving as a positive control (Figure 1B,C), validating correct genomic integration. When we crossed Ntsr1^Flp+/-^ to C57BL/6J, we obtained the expected Mendelian ratio of progeny: 50.7% Flp^+^ and 49.3% Flp^-^ (P value = 0.90, 34 *Flp*^+^ and 33 *Flp*^-^; Chi squared = 0.015 with 1 degree of freedom).

**Figure 1:**
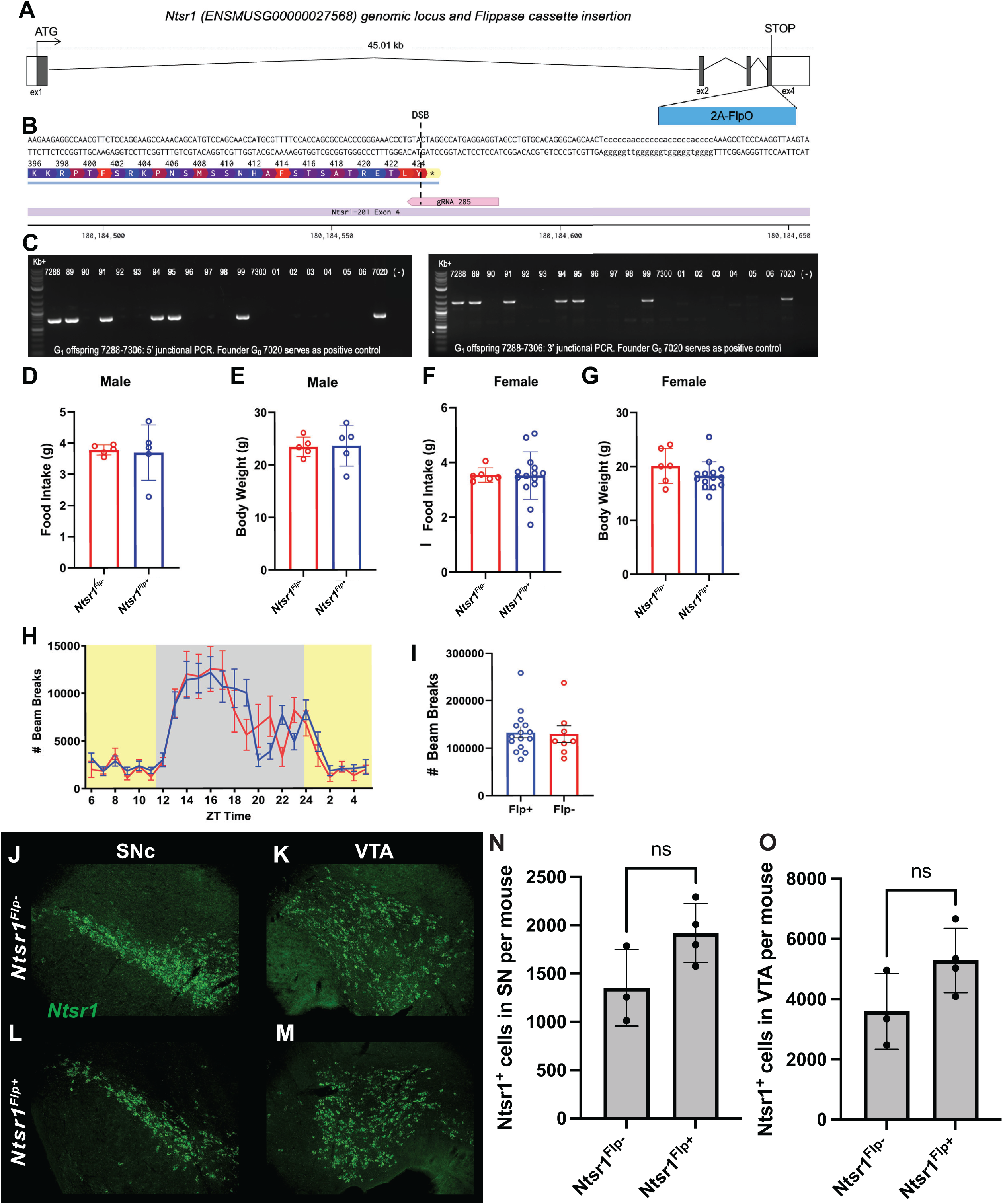
Generation and Physiological Characterization of Ntsr1^Flp^ mice. (A) Genomic locus of *Ntsr1* showing coding (filled boxes) exons and untranslated (white boxes) regions and insertion point of *2A-FlpO* cassette into the endogenous locus: overview (A) and precise location in exon 4 (B) with respect to the stop codon (*) with gRNA denoted in pink and the putative double stranded break generated by Cas9 indicated as a dotted line. (C) Genotyping of G1 offspring for the correctly targeted allele with a 5’ and 3’ junctional PCR with one external and one internal primer. Full Sanger sequencing of the 2A-FlpO cassette, flanking homology region and insertion junctions was completed to confirm the edited allele was intact. (D–G) Daily food intake and body weight in male and female Flp^−^ and Flp^+^ mice. (H) Circadian locomotor activity assessed by infrared beam breaks across Zeitgeber time (ZT). Shaded regions indicate the dark phase. (I) Total locomotor activity as cumulative beam breaks. Data are shown as mean ± SEM. Sample sizes: males, n = 5 pairs; females, n = 6 pairs; locomotor activity, Flp^+^ (n = 15) and Flp^−^ (n = 8). (J-M) Representative images of SN and VTA RNAscope staining for *Ntsr1*. (J) The SN and (K) the VTA of a *Flp*^-^ mouse and (L) the SN and (M) the VTA of a *Flp*^+^ mouse showing *Ntsr1* expression in green. (N) Quantification of Ntsr1^+^ expressing cells in SN in *Flp*^+^ and *Flp*^-^ and *Flp*^-^ mice. (O) Quantification of *Ntsr1*^+^ cells in VTA per *Flp*^+^ and *Flp*^-^ mice. (O) Quantification of *Ntsr1*^+^ cells in VTA per *Flp*^+^ and *Flp*^-^ mice.

To determine whether Flp insertion altered baseline physiology, we assessed metabolic and behavioral parameters in Flp⁺ and Flp⁻ littermates. In male mice (n = 5 per group), average daily food intake did not differ between genotypes (p = 0.84, unpaired two-tailed t-test; Figure 1D). Similarly, male body weight was comparable between groups (p = 0.91, unpaired two-tailed t-test; Figure 1E). In female mice (n = 6 per group), food intake was also unchanged between Flp⁺ and Flp⁻ animals (p = 0.95, unpaired two-tailed t-test; Figure 1F). Female body weight showed no significant effect of genotype (p = 0.20, unpaired two-tailed t-test; Figure 1G).

We next evaluated circadian locomotor activity using infrared beam-break monitoring across the light–dark cycle in the home-cage. Two-way repeated-measures ANOVA revealed a significant main effect of Zeitgeber time (ZT) (p < 0.0001, F(6.47, 135.9) = 21.67; *Flp*, n=15; *Flp*, n=8, pooled sexes), reflecting normal nocturnal activity, but no main effect of genotype (p = 0.85, F(1,21) = 0.039) and no time × genotype interaction (p = 0.16, F(6.47, 135.9) = 1.556, Figure 1H). Total cumulative beam breaks over the recording period did not differ between genotypes *Flp* : 133,233; Flp : 129,277 (p = 0.85, unpaired two-tailed t-test, Figure 1I). We also used RNAScope to examine endogenous *Ntsr1* RNA expression RNA expression in the SN and VTA of *Ntsr1^Flp^*^⁺^ and *Ntsr1^Flp^*^⁻^ animals, with qualitatively similar anatomical distributions across genotypes (Figure 1J–M). Quantification revealed no significant genotype-dependent difference in the number of *Ntsr1+* cells in either SN or VTA (t test, p = 0.1139 for SN and p = 0.1341 for VTA; n=3 *Ntsr1^Flp-^* and n=4 *Ntsr1^Flp+^*; Figure 1J-O). Collectively, these data demonstrate that targeted insertion of the *2A-FlpO* cassette into the *Ntsr1* locus does not produce detectable alterations in baseline food intake, body weight, or locomotor activity, supporting the physiological suitability of this line for intersectional genetic studies.

### Intracranial AAV injection to assess Flp-dependent mCherry expression in the SN and VTA of Ntsr1^Flp^ mice

To validate Flp-dependent recombination in the midbrain, we injected AAV9-EF1α-fDIO-mCherry bilaterally in the SN of *Ntsr1^Flp^* mice. Here, we observed mCherry expression mainly in tyrosine hydroxylase (TH)^+^ neurons, but less restricted than would be expected for a purely dopaminergic population. Quantification of mCherry⁺ neurons co-labeled with TH revealed that 65% were TH, whereas ∼35% lacked TH expression (Figure 2A-E).

**Figure 2.**
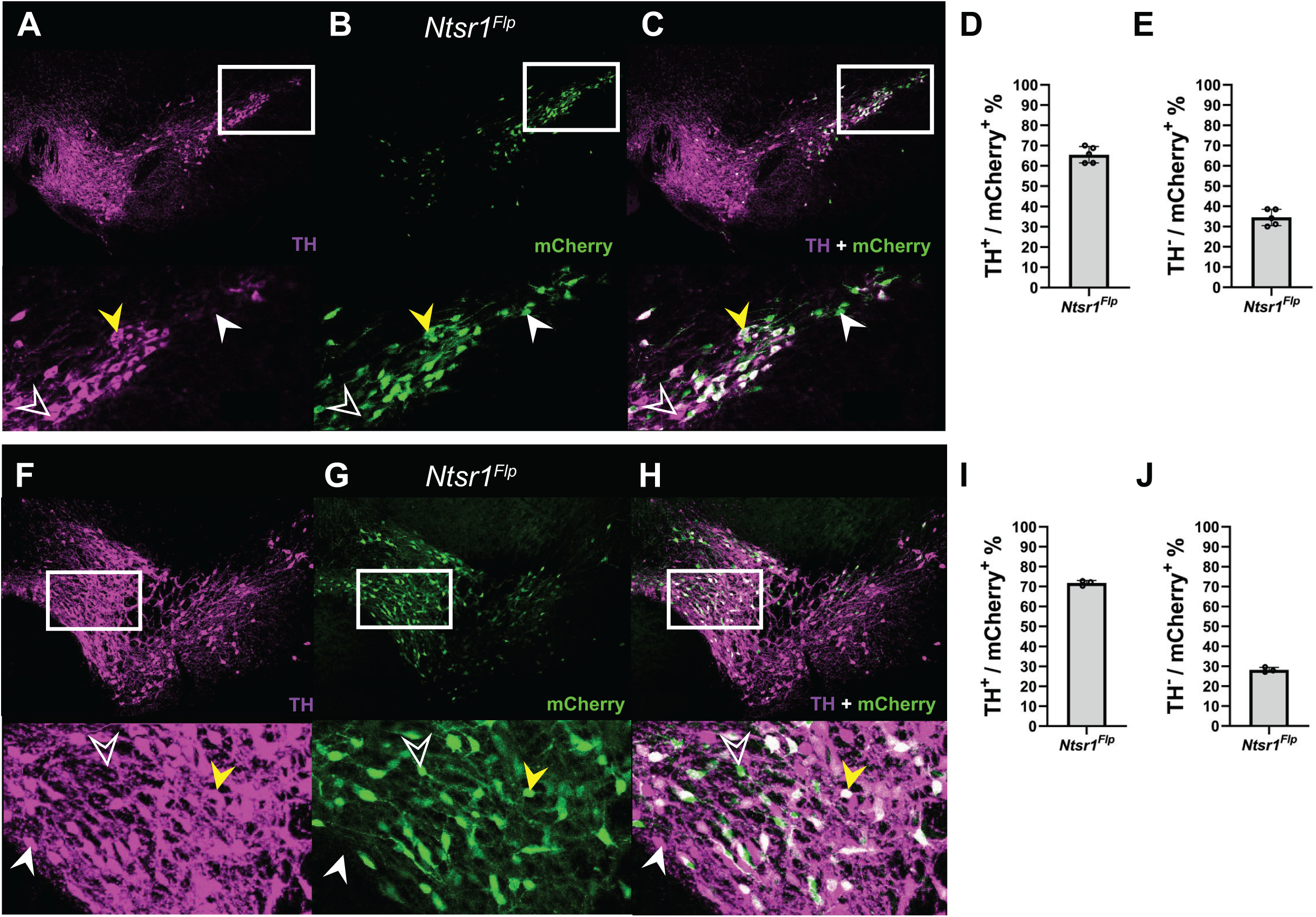
IC injection of AAV encoding Flp-dependent mCherry in the SN and VTA of Ntsr1^Flp^ mice. (A–C) TH immunostaining (magenta) and Flp-dependent mCherry expression (green) in the substantia nigra (SN) following bilateral injection of AAV9-EF1a-fDIO-mCherry (2.5 × 10^8^ vg per site). (A) TH labeling in the SN. (B) mCherry expression driven by Flp recombination. (C) Merged image showing colocalization of TH and mCherry (white). Insets show higher-magnification views of the boxed regions. Yellow arrowheads indicate TH^+^/mCherry^+^ neurons; open arrowheads indicate TH^+^ neurons lacking mCherry expression, and the white arrowheads show mCherry^-^ and TH^+^. (D,E) Percentage of TH^+^ neurons expressing mCherry (D) and percentage of mCherry+ neurons co-expressing TH (E) in the SN. (F–H) TH (magenta) and mCherry (green) labeling in the VTA. (F) TH labeling in the VTA. (G) Flp-dependent mCherry expression. (H) Merged image showing TH^+^/mCherry^+^ colocalization (white). Insets show boxed regions at higher magnification. (I,J) Percentage of TH^+^ neurons expressing mCherry (I) and percentage of mCherry^+^ neurons co-expressing TH (J) in the VTA. Data are mean ± SEM. SN: n = 5 mice; VTA: n = 3 mice.

We injected the same dose of AAV9-EF1α-fDIO-mCherry into the VTA and observed that ∼70% of mCherry neurons co-expressed TH and ∼30% were TH (Figure 2F–J). These results demonstrate efficient Flp-dependent recombination in midbrain Ntsr1 neurons and suggest that Ntsr1 expression may encompass both dopaminergic and non-dopaminergic populations in both the SN and VTA as assessed by TH antibody staining.

### Systemic AAV injection to assess Flp-dependent mNeonGreen expression in the brains of Ntsr1^Flp^ mice

To characterize the distribution of Flp recombinase activity in the brain of *Ntsr1^Flp^* mice, adult animals received systemic injections of PHP.eB-CAG-fDIO-mNeonGreen (3E11 vg, n=3). Systemic delivery of the reporter resulted in mNeonGreen expression in several locations in the brains of *Ntsr1^Flp^* mice (Figure 3). In addition to striatal labeling, we also observed thalamic labeling (Figure 3A). Even more labeled cells were present in the mediodorsal nucleus of the thalamus (Figure 3B). Dense mNeonGreen-positive axonal projections were observed, likely extending from the midbrain, consistent with known nigrostriatal projections (Figure 3C). Cortical expression of the Flp-reporter was surprisingly weak, as there were a handful of labeled cortical neurons per section (Figure 3D). One of the only other regions where we noted staining was in a sparsely labeled population of cells visible in the diagonal band (Figure 3E). To determine whether Ntsr1-expressing neurons overlapped with dopaminergic populations, sections containing the VTA and SN were immunostained for TH. Within the VTA and SN, mNeonGreen-positive cells exhibited substantial colocalization with TH, although TH^+^ neurons lacking mNeonGreen expression were also abundant in these regions (Figure 3F-G). As a negative control for the mNeonGreen viral leak, we injected Flp^-^ mice and their tissue exhibited little to no mNeonGreen fluorescence. Rare mNeonGreen-positive cells (approximately 3–4 cells per section) were observed in Flp^-^ brains, indicating minimal reporter leak expression (Figure 3H). None of these mNeonGreen^+^ cells co-expressed TH (Figure 3H, insets). Quantification revealed that approximately half of mNeonGreen-positive neurons in the SN were TH^+^ and half were TH^-^ (Figure. 3I). In contrast, the VTA contained a substantially smaller fraction of TH^-^;mNeonGreen^+^ neurons, while ∼70% mNeonGreen^+^ neurons in this region were TH^+^ (Fig. 3J).

**Figure 3.**
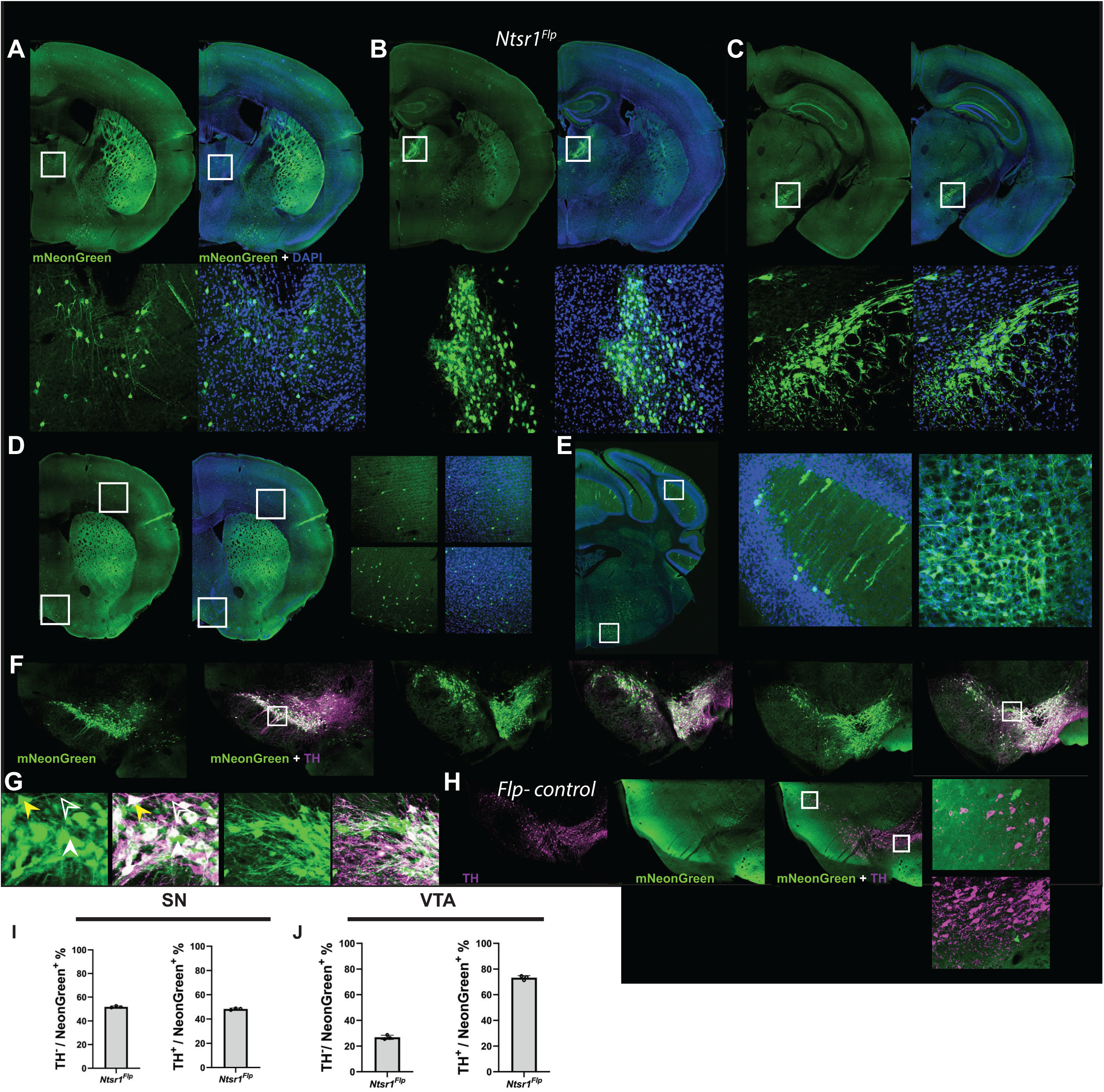
Characterization of systemic Flp-dependent mNeonGreen reporter expression in the brains of Ntsr1^Flp^ mice and overlap with dopaminergic neurons. (A–C) Representative coronal brain sections showing Flp-dependent mNeonGreen expression (green) in *Ntsr1^Flp^* mice. Boxed regions are shown at higher magnification below each panel. DAPI (blue) labels cell nuclei. (D,E) Additional representative brain regions illustrating the distribution of mNeonGreen-positive neurons and axonal projections. Insets show higher-magnification views of boxed regions. (F) Representative images of the SN and VTA immunostained for TH (magenta) together with mNeonGreen (green). (G) Higher-magnification views of the insets from SN. Arrowheads indicate representative neurons positive for both mNeonGreen and TH (yellow arrowheads) or cells expressing only TH (open arrowhead) or expressing mNeonGreen only (white arrowhead). (H) Representative images from Flp-negative control tissue demonstrating several mNeonGreen^+^ cells. Insets show that these cells do not co-stain for TH. (I) Quantification of overlap between mNeonGreen and TH labeling in the SN, expressed as the percentage of TH-positive neurons that were mNeonGreen-positive and the percentage of mNeonGreen-positive neurons that were TH-positive. (J) Quantification of overlap between mNeonGreen and TH labeling in the ventral tegmental area. Data are presented as mean ± SEM.

### Systemic AAV injection to assess Cre- and Flp-dependent mCherry expression in the SN and VTA of dual-recombinase mice

Given the unexpectedly large TH^−^ fraction among Flp-recombined cells, we increased targeting stringency by requiring coincident Cre and Flp-dependent activity using a systemically delivered Cre-on/Flp-on reporter. We employed a dual-recombinase Cre-on/Flp-on (‘Con/Fon’) construct to express mCherry packaged in blood-brain barrier crossing PHP.eB capsid (Figure 4).^27,33–35^ PHP.eB-nEF-Con/Fon-TVA-mCherry was delivered via retro-orbital (RO) injection into complementary mouse recombinase configurations utilizing*Dat^Flp^*,^36^ *Dat^Cre^*,^37^ *Ntsr1^Cre^*,^22^ and *Ntsr1^Flp^* (created this study): Dat^Cre^;Ntsr1^Flp^ and Ntsr1^Cre^;Dat^Flp^.

**Figure 4.**
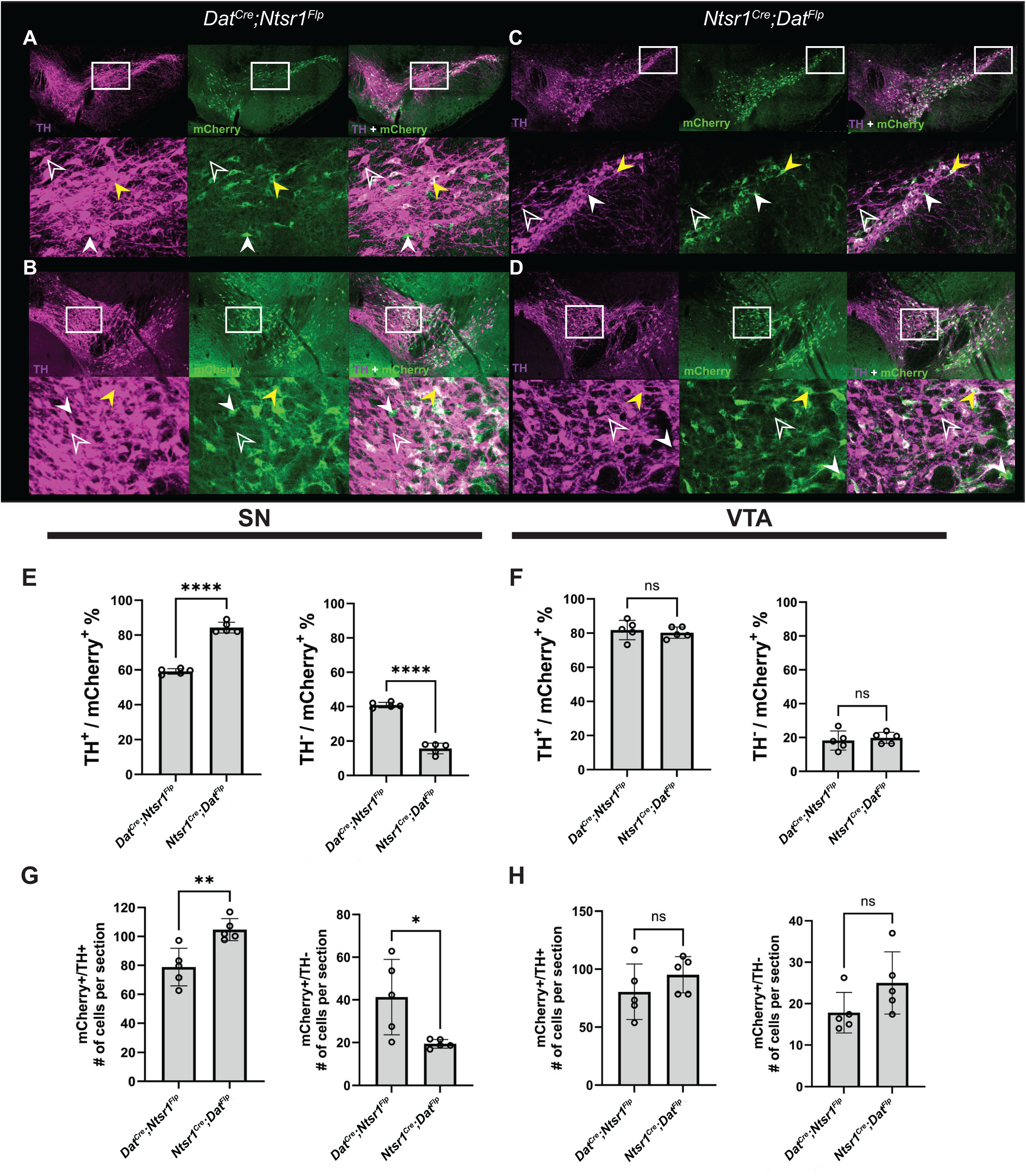
Configuration-dependent differences in dopaminergic specificity with systemic AAV expressing dual-recombinase-dependent mCherry. (A–C) TH immunostaining (magenta) and intersectional mCherry expression (green) in the SN of Dat^Cre^;Ntsr1^Flp^ mice following systemic delivery of PHP.eB-nEF-Con/Fon-TVA-mCherry. (A) TH labeling. (B) mCherry expression. (C) Merged image showing colocalization. Insets show higher magnification. Arrowheads indicate TH^+^,mCherry+, and TH⁺/mCherry⁺ neurons; Yellow arrowheads indicate TH^+^/mCherry^+^ neurons; open arrowheads indicate TH^+^ neurons lacking mCherry expression, and the white arrowheads show mCherry^-^ and TH^+^ . (D,E) Quantification in the SN of the percentage of TH⁺ neurons expressing mCherry (D) and mCherry⁺ neurons co-expressing TH^-^ (E). (F–H) TH and mCherry labeling in the VTA of Dat^Cre^;Ntsr1^Flp^ mice. (I,J) Quantification in the VTA of TH⁺/mCherry⁺ and TH⁻/mCherry⁺ neurons in fractions. (K–M) TH and mCherry labeling in the SN of Ntsr1^Cre^;Dat^Flp^ mice. (N,O) Quantification in the SN comparing recombinase configurations.(P–R) TH and mCherry labeling in the VTA of Ntsr1^Cre^;Dat^Flp^ mice. (S,T) Absolute numbers of TH⁺/mCherry⁺ neurons per section in SN and VTA. Data are mean ± SEM; n = 5 mice per genotype. Statistical comparisons were performed using unpaired two-tailed t-tests.

In the SN of *Dat^Cre^;Ntsr1^Flp^* mice, TH immunostaining and mCherry expression in the SN showed considerable but not overwhelming overlap (Figure 3A,C). Quantification of the percentage of TH neurons co-expressing mCherry was 59.1% while the TH^-^ expressing mCherry was 40.9% (p=<0.0001, unpaired two-tailed t-test, Figure 4,E). In contrast, the reciprocal *Ntsr1^Cre^*;*Dat^Flp^* configuration yielded significantly greater dopaminergic specificity in the SN, with 84.3% of mCherry⁺ neurons co-expressing TH and 15.7% TH⁻ (p < 0.0001; n = 5 per group; Figure 4E).

Absolute cell counts mirrored these proportional differences. In the SN, *Dat^Cre^;Ntsr1^Flp^* mice exhibited an average of 79 TH⁺/mCherry⁺ neurons and 40 TH⁻/mCherry⁺ neurons per section, whereas *Ntsr1^Cre^;Dat^Flp^*mice showed 105 TH⁺/mCherry⁺ neurons and 20 TH⁻/mCherry⁺ neurons (Figure 4G). Together, these data indicate that recombinase configuration significantly influences dopaminergic enrichment in the SN.

In the VTA, however, recombinase configuration did not significantly alter dopaminergic targeting specificity (Figure 4F, H). *Dat^Cre^;Ntsr1^Flp^* and *Ntsr1^Cre^;Dat^Flp^* mice exhibited comparable proportions of TH⁺/mCherry⁺ neurons (80.2% vs. 81.8%, respectively; p = 0.61) and TH⁻/mCherry⁺ neurons (19.8% vs. 18.2%; p = 0.61). Absolute counts were similarly comparable, with approximately 80 vs. 95 TH⁺/mCherry⁺ neurons and 19 vs. 25 TH⁻/mCherry⁺ neurons per section for *Dat^Cre^;Ntsr1^Flp^* and *Ntsr1^Cre^;Dat^Flp^* mice, respectively (Figure 4H). Together, these findings demonstrate that recombinase configuration enhances dopaminergic targeting specificity in the SN but does not significantly alter targeting efficiency in the VTA, revealing region-dependent differences in intersectional recombination performance.

### Systemic AAV injection to assess Cre- and Flp-dependent mCherry expression in the SN and VTA of cis-gene dual recombinase controls

To determine the upper bound of recombinase-dependent targeting efficiency, we examined “cis-gene” dual recombinase configurations in which both Cre and Flp are expressed from the same endogenous locus. We intercrossed *Dat^Cre^*with *Dat^Flp^*; similarly, we intercrossed *Ntsr1^Cre^*with *Ntsr1^Flp^*. Mice were injected RO with PHP.eB Cre-on/Flp-on mCherry reporter under the same conditions as above.

In the SN, *Dat^Cre^;Dat^Flp^* mice exhibited very high colocalization between mCherry and TH (Figure 5A). Approximately ∼90% of mCherry neurons were TH, with a correspondingly low TH fraction (Figure 5E). In contrast, *Ntsr1^Cre^;Ntsr1^Flp^* (Figure 5C) mice showed significantly reduced dopaminergic specificity in the SN (∼70% TH /mCherry), accompanied by a substantial increase in TH^+^/mCherry neurons at around 89.6% vs 71.25% (p<0.0001 unpaired two-tailed t-test, Figure 5F). Correspondingly, the proportion of TH /mCherry neurons was significantly elevated in *Ntsr1^Cre^*;*Ntsr1^Flp^* mice relative to *Dat^Cre^*;*Dat^Flp^* cis mice 28.8% vs 10.4% (p<0.0001, unpaired two-tailed t-test, Figure 5E). Analysis of absolute cell counts in the SN revealed a significantly higher number of TH /mCherry neurons in *Dat^Cre^;Dat^Flp^* mice compared to *Ntsr1^Cre^;Ntsr1^Flp^* mice 121.1 vs 35.1 cells (p<0.0001, unpaired two-tailed t- test, Figure 5G). In contrast, TH /mCherry cell counts did not significantly differ between genotypes 12.4 vs 14.27 cells (p=0.37, unpaired two-tailed t-test, Figure 5G).

**Figure 5.**
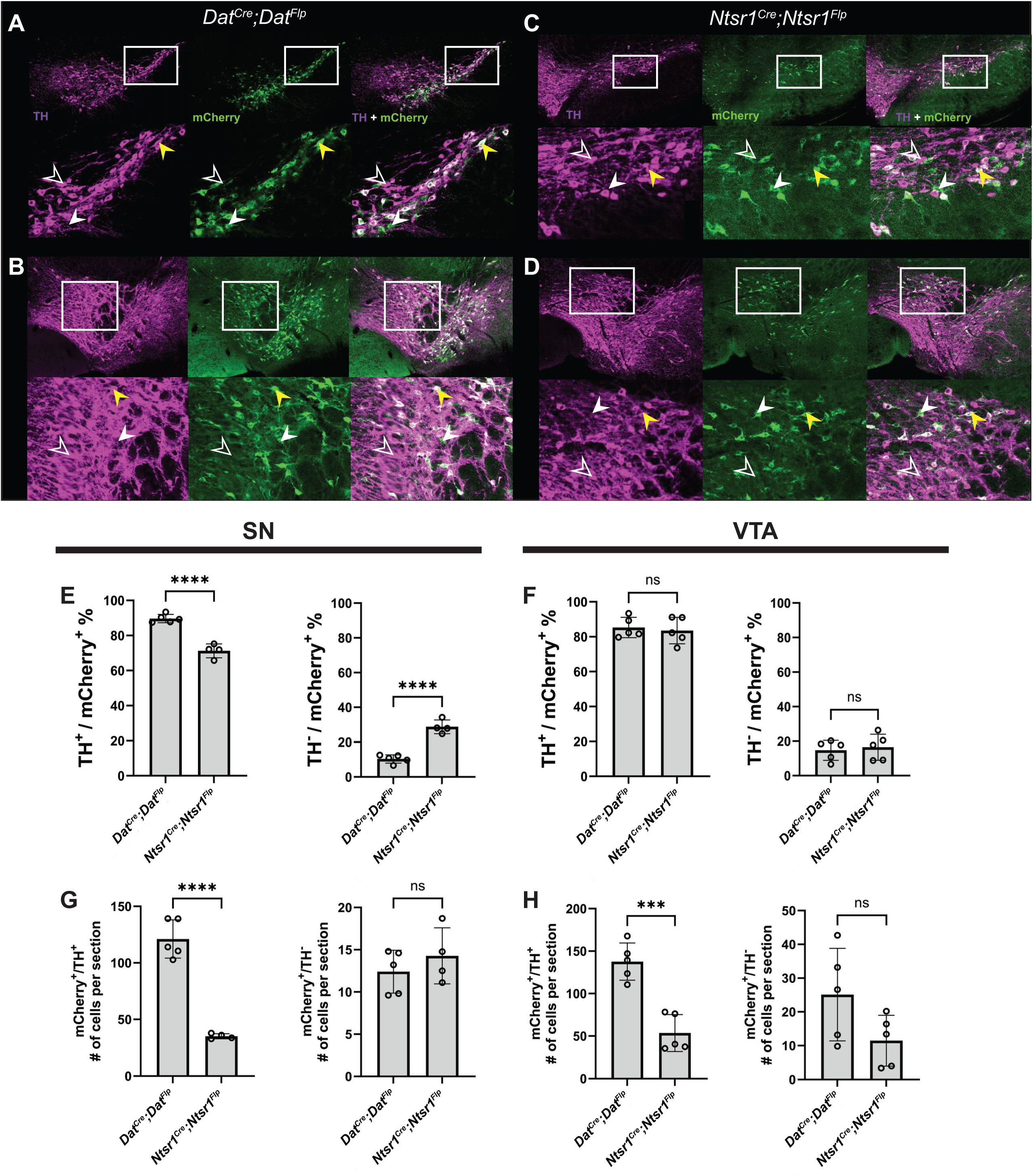
Cis-gene dual recombinase controls define maximal dopaminergic targeting specificity with systemic AAV expressing dual recombinase-dependent mCherry. (A–C) TH immunostaining (magenta) and intersectional mCherry expression (green) in the SN of *Dat^Cre^;Dat^Flp^* mice. Insets show higher-magnification views of the boxed regions. Yellow arrowheads indicate TH^+^/mCherry^+^ neurons; open arrowheads indicate TH^+^ neurons lacking mCherry expression, and the white arrowheads show mCherry^-^ and TH^+^. (D–F) TH and mCherry labeling in the VTA of *Dat^Cre^;Dat^Flp^* mice. (G–I) TH and mCherry labeling in the SN of Ntsr1^Cre^;Ntsr1^Flp^ mice. (J–L) TH and mCherry labeling in the VTA of *Ntsr1^Cre^;Ntsr1^Flp^* mice. (M,N) Quantification in the SN of TH⁺/mCherry⁺ and TH⁻/mCherry⁺ fractions. (O,P) Quantification in the VTA. (Q,R) Absolute TH⁺/mCherry⁺ and TH⁻/mCherry⁺ cell counts in SN. (S,T) Absolute counts in VTA. Data are mean ± SEM. Statistical comparisons were performed using unpaired two-tailed t-tests.

In the VTA, the percent colocalization between TH and mCherry did not significantly differ between cis configurations (ns, unpaired two-tailed t-test, Figure 5B,D, F). The proportion of TH⁺/mCherry⁺ neurons did not significantly differ between cis configurations (Dat: 85.3% vs Ntsr1: 83.5%; p=0.70; Figure 5F). Similarly, the TH /mCherry fraction was comparable between groups (p=0.70, unpaired two-tailed t-test, Figure 5F). However, absolute counts of TH /mCherry neurons were significantly higher in *Dat^Cre^;Dat^Flp^* mice compared to *Ntsr1^Cre^;Ntsr1^Flp^* mice 137.6 vs 53.5 cells; (p=0.0003, unpaired two-tailed t-test, Figure 5H). Absolute counts of TH^-^ /mCherry neurons remained higher in *Dat^Cre^;Dat^Flp^* mice at 25.1 vs 11.5 cells. Absolute TH /mCherry counts in the VTA did not reach statistical significance (p=0.09, unpaired two-tailed t-test, Figure 5H), consistent with stronger dopaminergic restriction in this configuration. Together, these results demonstrate that when both recombinases are driven from the *Dat* locus, intersectional recombination approaches a maximal dopaminergic ceiling. The persistence of TH /mCherry neurons in *Ntsr1^Cre^;Ntsr1^Flp^*mice likely reflects the intrinsic cellular heterogeneity of *Ntsr1* expression rather than incomplete recombinase efficiency.

As a test for leakiness of the virus, we examined expression of mCherry delivered from PHP.eB-nEF-Con/Fon-TVA-mCherry in the following negative control genotypes: *Ntsr1^Cre-^*;*Ntsr1^Flp+^*or *Ntsr1^Cre-^*;*Dat^Flp+^*. Neither of these mice showed any expression in the ventral midbrain from systemic AAV (Supplemental Figure 1A-L). We also looked for mCherry expression in the forebrain of all the dual recombinase genotypes, but we did not observe any expression, suggesting minimal recombinase-independent reporter activation (Supplemental Figure 1M-X).

Lastly, to determine if TH expression was missing a portion of genuine dopaminergic neurons, we co-stained with TH and DAT antibodies. We observed a small subset of TH⁻/DAT⁺/mCherry⁺ neurons in both the SN and VTA (Supplementary Figure 2). From cell counts of n=3 mice of the following genotypes *Dat^Cre^*;*Dat^Flp^*, *Dat^Cre^*;*Ntsr1^Flp^*, *Ntsr1^Cre^*;*Dat^Flp^*, we estimate that there were 4.4% TH^-^DAT^+^mCherry^+^ cells and that using TH antibody staining only causes an underestimate of the fraction of dopaminergic co-localization with mCherry.

### Intracranial AAV injection of Cre- and Flp-dependent taCaspase-3 induces dopaminergic ablation in the SN of dual-recombinase mice

To determine whether intersectional recombination could be leveraged for selective cell ablation, we designed and employed a Cre- and Flp-dependent Con/Fon taCasp-3 TEVp construct.^31,32^ The design of the EF1A-Con/Fon-taCasp3 construct enabling Cre and Flp-dependent reorientation of the active caspase-3 cassette is illustrated schematically pre- (Figure 6A) and post-recombinase activity (Figure 6B). To test the functional expression of this construct, DJ/9-EF1A-Con/Fon-taCasp3 was delivered bilaterally to the SN of Cre^−/^Flp^−^ controls, single-recombinase mice (*Dat^Cre^*, *Ntsr1^Cre^*, *Dat^Flp^*, or *Ntsr1^Flp^*), and Cre^+^/Flp^+^ mice (n=2 *Dat^Cre^*;*Ntsr1^Flp^*and n=1 *Ntsr1^Cre^*;*Dat^Flp^*). Quantification of dopaminergic neuron survival revealed statistical differences across recombinase conditions (Figure 6C). Representative TH immunostaining images of the SN are shown for Cre^−^/Flp^−^ controls (Figure 6D), Cre^+^/Flp^−^ mice (Figure 6E), Cre^−^/Flp^+^ mice (Figure 6F), and Cre^+^/Flp^+^ mice (Figure 6G). TH immunostaining revealed preserved dopaminergic neuron density in Cre^−^/Flp^−^ controls and in both single-recombinase conditions (Figure 6D–F). In contrast, Cre^+^/Flp^+^ mice exhibited a marked reduction in TH⁺ neurons within the SN (Figure 6G), consistent with dual recombinase-dependent caspase activation. Quantification confirmed a significant effect of recombinase condition on TH⁺ cell number relative to the double-positive recombinase condition (p value = 0.0001, one-way ANOVA with Tukey’s post hoc test; Figure 5C). Cre^+^;Flp^+^ mice displayed approximately 50 TH⁺ cells per section of SN compared to ∼220 cells per section in Cre^−^/Flp^−^ controls (p = 0.0002, Tukey Multiple Comparisons, Figure 5C). Similar results were obtained for single-recombinase controls ∼220 cells per section (p = 0.0002, Tukey Multiple Comparisons, Figure 6C). These results demonstrate that dual-recombinase activation of taCasp3 is sufficient to drive robust, cell-type–restricted ablation of midbrain dopaminergic neurons in vivo.

**Figure 6.**
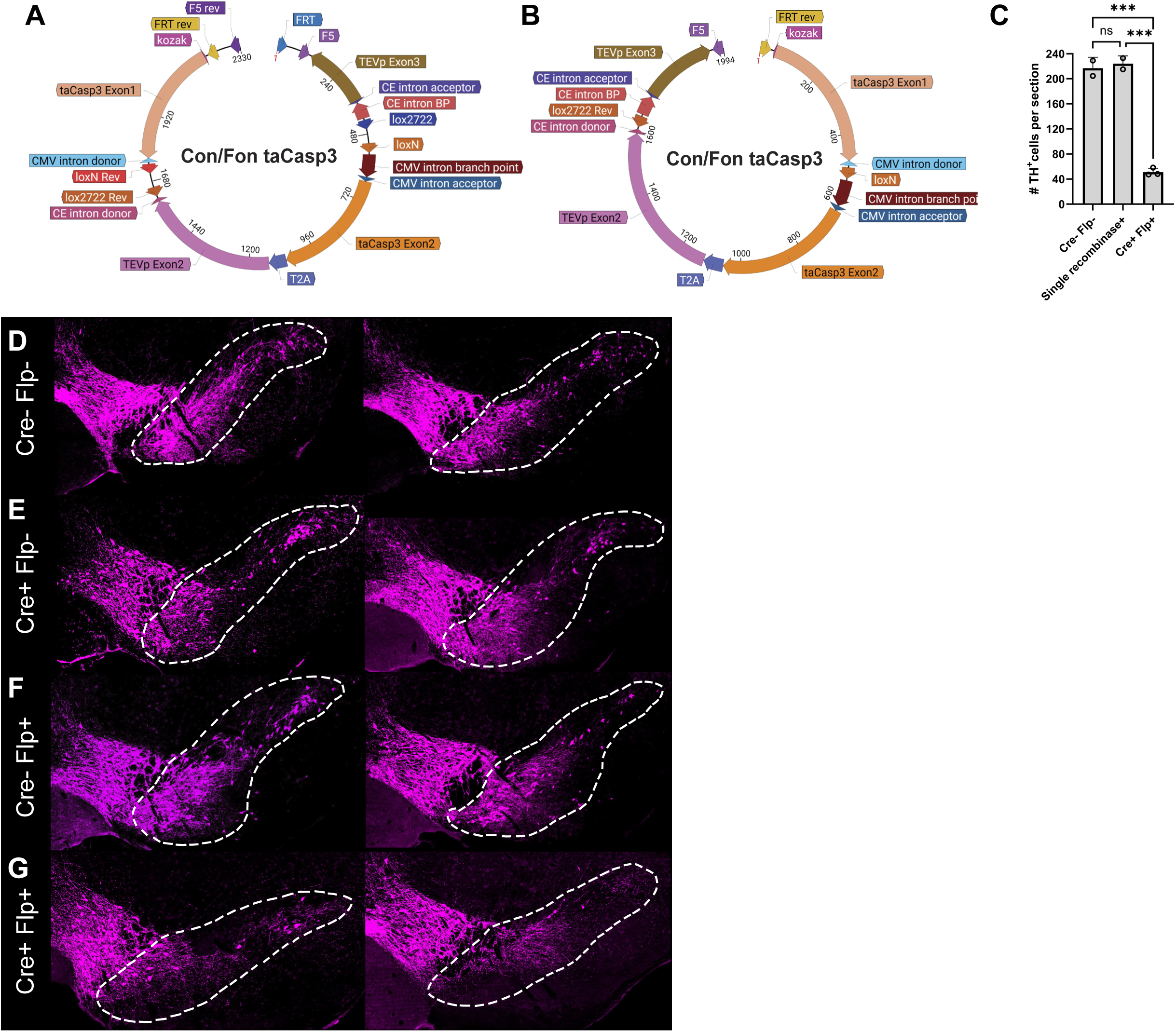
Intersectional recombination drives taCasp3-TEVP mediated ablation of TH+ neurons using direct injection of AAV. (A) Schematic of the Con/Fon taCasp3 construct prior to recombination. The cassette contains inverted taCasp3 exons flanked by loxP/lox2722 and FRT sites under control of the EF1A promoter. (B) Post-recombination configuration following Cre- and Flp-mediated recombination, resulting in correctly oriented taCasp3 expression.(C–G) TH immunostaining in the midbrain across recombinase conditions following DJ9-EF1A-Con/Fon-taCasp3 delivery. (C) Cre^−^/Flp^−^ control. (D–F) Single recombinase^+^ controls. (G) Cre^+^/Flp^+^ condition demonstrating marked reduction of TH^+^ neurons consistent with caspase-mediated ablation. (C) Quantification of TH^+^ neurons per section across recombinase conditions through the use of a One-way ANOVA. Data are shown as mean ± SEM. Cre^−^/Flp^−^ (n = 2 mice), Cre^+^/Flp^+^ (n = 3 mice), single recombinase^+^ (n = 2 mice total). Statistical comparisons were performed using one-way ANOVA followed by Tukey’s multiple comparisons test. Statistically significant differences are indicated as follows: ns, not significant; ∗p < 0.05; ∗∗p < 0.01; ∗∗∗p < 0.001.

## Discussion

Recent studies suggest that distinct sites of Ntsr1-expressing neurons in the brain mediate different aspects of the wide swath of physiology and behavior attributed to Nts signaling. Mapping the sites and identities of Ntsr1 neurons and the effects they mediate can reveal how they contribute to normal physiology as well as how and where they might be leveraged to address dysfunction. For example, Nts signaling has been shown to increase feeding via some brain areas while it decrease it via others.^23,38–41^ Moreover, transcriptomics are revealing that there are subsets of Ntsr1 neurons even within the same brain region which may have distinct functions. Hence, it is imperative to understand how site- and molecularly-specified populations of Ntsr1 neurons mediate physiology and behavior, but this requires tools capable of distinguishing them.

To that end, in this study, we developed and validated a knock-in *Ntsr1^Flp^* recombinase line that enables selective and intersectionally compatible genetic access to Ntsr1-expressing neurons. *Ntsr1^Flp^*preserves endogenous Ntsr1 neuronal function, supports both local and systemic AAV strategies, and reveals previously underappreciated cellular heterogeneity within midbrain Ntsr1 populations. Importantly, insertion of the *P2A-FlpO* cassette did not perturb baseline physiology: *Ntsr1^Flp^* mice were born at expected Mendelian ratios and exhibited normal body weight, food intake, and locomotor activity. Given the established role of Ntsr1 signaling in energy balance, this physiological neutrality is critical, indicating that the knock-in allele does not introduce detectable haploinsufficiency or behavioral confounds. Together, these findings establish *Ntsr1^Flp^*as a robust and physiologically suitable tool for dissecting midbrain circuits involved in feeding, drinking, motivation, and metabolic regulation. Importantly, *Ntsr1^Flp^* enables experimental strategies that are not achievable using *Ntsr1^Cre^* alone.^21^ By combining *Ntsr1^Flp^* with dopaminergic, projection-specific,^35^ or enhancer-driven Cre lines,^42^ investigators can isolate Ntsr1⁺ dopaminergic neurons from Ntsr1⁺ non-dopaminergic populations or conversely, restrict manipulations to non- dopaminergic Ntsr1 subsets through Boolean exclusion logic. This flexibility permits causal testing of molecularly defined subpopulations within circuits regulating feeding,^10^ motivation, and metabolic control,^23^ and expands the experimental repertoire beyond what is possible with single recombinase strategies, facilitating systemic rather than surgical delivery of AAVs.

Using intracranial delivery of a Flp-dependent reporter, we observed that *Ntsr1^Flp^* activity expression in the SN and VTA encompasses both TH^-^ and TH^+^ neurons. The substantial TH⁻ fraction was unexpected and challenges the assumption that Ntsr1 selectively marks dopaminergic neurons in the ventral midbrain. Instead, our data support a model in which Ntsr1 defines a molecularly heterogeneous population within both the SN and VTA that encompasses both dopaminergic and non-dopaminergic neurons. This observation has important implications for prior studies that relied on *Ntsr1^Cre^*or Ntsr1-based viral strategies without neurotransmitter restriction, as behavioral and physiological manipulations previously attributed to dopaminergic Ntsr1 neurons may, in part, reflect contributions from non-dopaminergic Ntsr1 populations. These cells may represent GABAergic or glutamatergic neurons participating in local nigral or mesencephalic circuits, although their precise identity remains to be determined. Intersectional strategies enabled by *Ntsr1^Flp^* provide a direct means to resolve these contributions.

Several lines of evidence argue against viral inefficiency or leakage as primary explanations for the TH⁻/mCherry⁺ population. Intracranial delivery produces a high multiplicity of infection, maximizing recombination across Flp-expressing neurons, and similar TH⁺/TH⁻ proportions were observed in both SN and VTA, arguing against stochastic under-infection. Similar results were obtained with systemic injection of fDIO systemic Con/Fon reporter expression was restricted to the midbrain despite widespread PHP.eB distribution,^34^ and negative controls containing only a single recombinase expression (*Ntsr1^Cre-^*/*Ntsr1^Flp+^*or *Ntsr1^Cre-^*/*Dat^Flp+^*) showed no reporter expression (Supplementary Figure 1), excluding recombinase-independent activation of the viral construct.

We observed an configuration-dependent difference in targeting specificity within the SN, where the *Ntsr1^Cre^;Dat^Flp^* configuration yielded higher dopaminergic specificity than the reciprocal *Dat^Cre^;Ntsr1^Flp^*configuration. Since these experiments utilized systemic AAV delivery in adult mice, this asymmetry cannot be attributed to transient developmental expression of the drivers. Instead, we hypothesize that this difference arises from the relative expression levels and catalytic efficiencies of the two recombinases. Cre recombinase has been reported to exhibit higher catalytic efficiency than FlpO under certain conditions.^43–45^ In the *Ntsr1^Cre^;Dat^Flp^* configuration, the ’weaker’ or more stringently regulated *Dat* driver (expressing FlpO) may act as a more effective secondary gate, ensuring that recombination only occurs in cells with high, definitive dopaminergic character. In contrast, in the *Dat^Cre^;Ntsr1^Flp^*configuration, the highly efficient Cre-recombinase may facilitate recombination in cells where *Ntsr1* (driving FlpO) is expressed at lower or non-specific levels. These results suggest that for intersectional targeting of heterogeneous populations, the choice of which driver controls which recombinase is a variable that must be empirically validated to optimize specificity

Importantly, cis-gene *Dat^Cre^;Dat^Flp^* controls achieved ∼90% TH-mCherry colocalization using the same Con/Fon strategy, demonstrating high recombination efficiency within bona fide dopaminergic neurons. Prior reports have described *Dat^Cre^* as ∼96% specific and *Dat^Flp^*as ∼97% specific following local viral delivery,^46^ with *Dat^Flp^* specificity modestly reduced (∼92%) when crossed to Cre-dependent reporters rather than using viral strategies.^36^ These benchmarks indicate that dual-recombinase targeting in dopaminergic neurons can approach near-ceiling specificity under optimal conditions. The persistence of a substantially larger TH⁻ fraction in Ntsr1-based configurations therefore exceeds what would be expected from recombinase inefficiency alone and instead supports intrinsic heterogeneity of Ntsr1 expression within the ventral midbrain. Future experiments employing mixed injections of Cre-dependent or Flp-dependent cis-cross configurations could further resolve potential configuration-dependent recombination asymmetries.

Incomplete detection of TH protein may contribute modestly to the observed TH⁻ fraction. TH expression varies across dopaminergic subtypes and physiological states, and antibody-based detection may fail to identify neurons with low or transient TH levels [32]. TH exhibits diurnal variation, with lower expression during the light phase [33,34], and animals in this study were perfused during the day. Consistent with this possibility, a small subset of mCherry⁺ neurons co-expressed DAT but not TH (Supplementary Figure 2), suggesting that a minority of mCherry labeled neurons may represent dopaminergic lineage cells with TH expression below detection threshold at the time of analysis. However, the systematic configuration-dependent differences observed across genotypes, together with high efficiency in cis-Dat controls, argue that incomplete TH detection cannot fully account for the magnitude of the TH⁻ population in either *Dat^Cre^;Ntsr1^Flp^* or *Ntsr1^Cre^;Dat^FLP^*VTA or SN. Collectively, these findings support the conclusion that Ntsr1 expression defines a genuinely heterogeneous neuronal population within the SN and VTA, comprising both dopaminergic and non-dopaminergic neurons.

Finally, we demonstrate that Ntsr1^Flp^ supports functional and/or behavioral studies through intersectional caspase-based ablation. To our knowledge, this represents the first demonstration of dual-recombinase–dependent ablation of neurons using a Con/Fon taCaspase-3 strategy in vivo. This functional validation extends the utility of the *Ntsr1^Flp^*line beyond reporter expression, establishing its compatibility with intersectional loss-of-function paradigms for causal circuit interrogation. Recombinase-dependent elimination of SN dopaminergic neurons confirms that this line enables efficient cell-autonomous manipulations beyond reporter expression. The compatibility of *Ntsr1^Flp^*with both systemic intersectional targeting and conditional ablation strategies positions it as a versatile tool for causal interrogation of defined midbrain circuits, particularly in concert with recently developed enhancer AAVs.^42^ In summary, the *Ntsr1^Flp^*mouse line provides robust and intersectionally compatible genetic access to Ntsr1-expressing neurons and uncovers unexpected cellular diversity within ventral midbrain Ntsr1 populations. By enabling both local and systemic dual-recombinase strategies, this resource facilitates refined dissection of dopaminergic-associated circuits and their non-dopaminergic counterparts.

## Methods

### Mouse husbandry, strains, and generation of Ntsr1-2A-FlpO mice

The experiments described herein were approved by the California State Polytechnic, Pomona Institutional Animal Care and Use Committee under protocols: 23.006, 23.009, and 22.020.

Mice used in experiments were maintained in a 12:12 light:dark cycle throughout all portions of study. Mice were housed in an Allentown NexGen Mouse 500 ventilated caging system (IVC). Zeitgeber Time (ZT) 0 was defined as lights on, whilst ZT 12 was lights off, by convention. Mouse cages were layered Sani-Chip bedding (Envigo, 7090), plastic or paper shelter structures, and cotton nestlets. The diet consisted of rodent chow 2018 Teklad 18% Protein Rodent Diet with caloric density of 3.1 kcal/g (Envigo). Ambient temperatures were maintained between 22-24°C with humidity between 20-45%.

*Ntsr1^Flp^* mice were generated by the Michigan State University Transgenic and Genome Editing Facility using CRISPR-Cas9-mediated HDR to insert an in-frame P2A-FlpO cassette at the C-terminus of the endogenous gene, immediately upstream of the native stop codon in exon 4, enabling bicistronic expression from the *Ntsr1* locus. Recombinant Cas9 protein was complexed with synthetic gRNA (Integrated DNA Technologies; N_20_ *PAM:* 5′-ACCTCCTCATGGCCTAGTAC *AGG*-3′) and microinjected into C57BL/6J zygotes together with a circular plasmid donor template (Twist Bioscience) carrying the 2A-FlpO cassette flanked by 500-bp 5′ and 3′ homology arms. Injected embryos were transferred into pseudopregnant recipients using standard methods. Founder mice were screened by junction PCR and validated by Sanger sequencing across both genomic–cassette junctions as well as the entire cassette and flanking homology regions. Germline transmission was confirmed in F1 offspring, which were subsequently backcrossed to C57BL/6J mice (The Jackson Laboratory, stock #000664) for three generations to stabilize the allele. Subsequent *NtsR1^2A-FlpO^*progeny were crossed with C57BL/6J mice to propagate the line and generate mice for studies. We performed SNP analysis using the MiniMUGA Background Analysis v2.3.1 (Transnetyx) on three mice (2 males and 1 female). All mice were a mixtures of C57BL/6J and C57BL/6NCrl and/or B6N-Tyr<c-Brd>/BrdCrCrl and/or C57BL/6NRj. The percent purity of the C57BL/6J strain was >96.5% for all three samples tested. The other mouse strains used in this study were *Dat^Cre^* (Jackson lab stock #020080), Dat^Flp^ (Jackson lab stock #035436), and *Ntsr1^Cre^*(Jackson labs stock #033365).

Locomotor activity was assessed using a photobeam break system (Columbus Instruments). Following acclimation to the chambers, spontaneous locomotor activity was continuously monitored for 72 hours using infrared beam-break detection. Activity counts were recorded and analyzed across both light and dark phases of the circadian cycle.

Body weight was measured using a precision digital balance. Daily food intake was calculated by subtracting the remaining food mass from the initial hopper weight, accounting for spillage where applicable.

### Viral Constructs and Administration

Recombinant adeno-associated viral vectors were used for Flp-dependent reporter expression. AAV9-EF1α-fDIO-mCherry (Addgene viral prep #114471-AAV9) was obtained from Addgene. The plasmid pAAV-EF1α-fDIO-mCherry was originally generated by Karl Deisseroth (Addgene plasmid #114471; RRID:Addgene_114471). The virus was used at 2.5 × 10 viral genomes (vg)/ul for per site for IC injections. The plasmid pAAV-nEF-Con/Fon-TVA-mCherry was obtained from Addgene (Addgene plasmid #131779; RRID:Addgene_131779). This plasmid was originally generated by Karl Deisseroth and was packaged as cargo in PHP.eB made in house at CPP. The pAAV-EF1a-Con/Fon taCasp3-T2A-TVEp_Modified:WPRE3 plasmid fully synthesized by Vectorbuilder (VB250909-1137abj) and packaged in AAVDJ/9^47^ in house. This AAV was injected at titer of 4.5 × 10^9^ vg in a total volume of 500nL per craniotomy site. AAVs were produced in-house using standard triple transfection and purification procedures following previously described protocols for AAV production.^29^

Male and female adult C57BL/6J *Ntsr1^Flp^* mice (8–10 weeks old; 22–28 g) were used for all experiments. All procedures were conducted in accordance with the NIH *Guide for the Care and Use of Laboratory Animals* and were approved by the IACUC. For preparation of IC, mice were anesthetized with isoflurane at an induction of 5%, maintenance at 2%, delivered in 0.5% oxygen (Fluriso, VetOne, #502017) and placed on a heated surgical pad to maintain body temperature. Anesthetized mice were secured in a stereotaxic apparatus (Kopf, Model 1900 Stereotaxic Alignment System). Following stabilization of the head, artificial tear ointment was placed onto the eyes and animals were injected subcutaneously (SQ) with 0.1mL 0.5mg/mL ketoprofen (Zoetis, Ketofen) and 0.5mL sterile saline (Hospira, 0.9% sodium chloride) for hydration during the procedure. The scalp was shaved and sterilized with alternating chlorhexidine and 70% isopropanol scrubs prior to incision. A midline scalp incision was made, and the skull was exposed. Skull leveling was performed both vertically and horizontally using a dual-probe leveling device (Kopf, Model 1905 Stereotaxic Alignment Indicator). After leveling, lambda was identified, and the sagittal suture was followed anteriorly until bregma was located. Coordinates relative to bregma were obtained using a stereotaxic frame with a digital micromanipulator. Bilateral craniotomies were drilled at coordinates relative to bregma for SN (ML: ±1.5 mm, AP: –3.6 mm, DV: –4.0 mm) and for the VTA (ML: ±0.65 mm, AP: –3.7 mm, DV: –4.0 mm) according to the *Allen Mouse Brain Atlas* (Allen Institute for Brain Science, 2011).

A glass needle filled with AAV9 was then lowered into each site, and 500 nL of viral solution (2.5 × 10 total) was delivered at a rate of 100 nL/min. The needle was left in place for at least 5 min post-injection to minimize backflow before being slowly withdrawn. After completion of viral delivery at both sites, the drill sites were sealed with bone wax (Surgical Specialties), and the scalp was closed using minimal tissue adhesive; VetBond (3M, No. 1469SB).

Bupivacaine (Hospira, Marcaine 0.5% 5mg/mL, PAA113011) was administered on the incision site to provide local analgesia. Ketoprofen (Zoetis, Ketofen 100mg/mL) was also administered subcutaneously at the conclusion of surgery. Following surgery, mice were placed under a heating lamp and monitored continuously until fully ambulatory. Once recovered, animals were returned to their home cages with food and water available *ad libitum*. Ketoprofen was administered subcutaneously once daily for 3 consecutive days. Two weeks post surgery, mice were then euthanized to test for transgene expression and/or cell ablation.

For RO injections, both male and female adult C57BL/6J *Ntsr1^Cre^*;*Dat^Flp^*, *Dat^Cre^*;*Ntsr1^Flp^*, *Dat^Cre^*;*Dat^Flp^*, and *Ntsr1^Cre^*;*Ntsr1^Flp^* mice aged 6–8 weeks old were used. Mice were anesthetized with isoflurane induction at 5%, maintenance at 2.5%, delivered in 0.5% oxygen for approximately five minutes. Injections were performed using a 30–33G insulin syringe with a fixed needle. Mice were positioned in lateral recumbency with the injection eye uppermost. The skin at the lateral canthus was gently retracted to expose the conjunctival margin without exerting pressure on the globe. The needle was introduced at the lateral canthus and advanced posteriorly along the orbital wall at approximately 30–45° toward the opposite mandible, passing behind the globe to access the retro-orbital venous sinus. Advancement continued until minimal resistance was encountered, indicating entry into the sinus vein. PhP.eB-nEF-Con/Fon TVA-mCherry was administered into the retro-orbital sinus at a concentration of 3 × 10^11^ vg in a total volume of 50 μL delivered slowly over ∼5 s. Following completion, the needle was withdrawn along the insertion path, and gentle pressure was applied with a sterile cotton-tipped applicator for 10–15 s to achieve hemostasis. The ocular surface of the injected eye received one drop of 0.5% proparacaine. Two weeks post surgery, the mice were then euthanized to test for transgene expression.

### Tissue Histology, Antibody Labeling, In Situ Hybridization, and Imaging

Transcardiac perfusion was performed on carbon dioxide euthanized mice by injecting 5-10 mL of phosphate buffer into the left ventricle followed by 2 mL of 4% PFA (Sigma), both made fresh prior to perfusion. Whole brain tissue was removed and stored in 4% PFA at -4C for at least 24 hours, then placed in 0.1M PBS. 50-micron coronal sections were obtained using a Leica VT1000S Vibratome (Leica Biosystems) and stored in 0.1M PBS at -4°C. Tissue samples were placed in glass well plates in PBST (0.5% Triton X-100 in PBS) to permeabilize the tissue for 10 minutes. Samples were then blocked in 5% goat serum for 10 minutes. Immunofluorescence antibody staining was performed using chicken polyclonal TH antibody (Aves, catalog TYH), rat DAT antibody (Millipore, MAB369), and DAPI (Invitrogen D1306). An Alexa Fluor 647-conjugated Goat Anti-Chicken IgG and an Alexa Fluor 488-conjugated Affinipure Goat anti-Rat (Invitrogen) where used to bind after the primary antibodies. Tissue samples were placed in a diluted TH primary antibody (Aves Lab), Rat Dopamine transport primary in goat serum at a dilution at 1: 500 and 1:250 overnight, and DAPI was used at 1:1000 then left overnight at 4C. Following overnight stain the tissue undergoes 5 minute washes three times in PBS. Secondary antibodies were used at 1:500 dilution for TH (Invitrogen AB_2535866) and DAT (InvitrogenAB_2534074) for approximately 1 hour then washed three times in PBS for 5 minutes each time. Sections were mounted using Immu-Mount (Epredia, #9990402) and visualized via confocal microscopy with a Nikon Eclipse Ti-E and NIS-Elements C software. Image processing and cell counting was performed with ImageJ (Fiji) using the Cell Counter plugin. Adjustments to images were performed using Fiji (formerly known as ImageJ). Confocal acquisition parameters were optimized independently for the SN and VTA to prevent detector saturation and maintain signals within the dynamic range. Due to regional differences in fluorescence intensity, laser power for the mCherry channel was reduced for VTA imaging relative to SN. Within each region, acquisition settings were held constant across all animals and sections. All images used for quantification were acquired below saturation.

For analysis, mCherry and TH^+^ cells were scored using binary criteria defined as signal clearly detectable above local background. Co-localization was defined as overlapping cellular signal in merged images. Quantification was based on cell counts and percent colocalization rather than fluorescence intensity measurements. Lookup table (LUT) ranges were standardized within each region across samples. For the mCherry channel, LUT values were set to 200–1250 for SN and 511–3700 for VTA. The TH channel was set to 511–3700 for both SN and VTA. For caspase-mediated ablation analysis of TH cells, the TH channel LUT was adjusted to 511–3700. LUT adjustments were applied for visualization and consistent thresholding within regions. For colocalization analysis, channels were pseudocolored (TH, magenta; TVA-mCherry, green), and merged images were generated. Cells expressing both TH and TVA-mCherry signals were manually counted. Quantification was performed bilaterally in both SN and VTA. Counts from left and right hemispheres were summed to generate a single value per section. Five rostro-caudal sections per animal were analyzed, and section values were combined to generate representative colocalization and ablation metrics for each genotype.

RNAscope for *Ntsr1* were performed as previously described.^21^ Quantification of RNAscope data was performed across four sections per mouse spanning from Bregma -3.64 to -2.92 to capture both VTA and SNc obtained from n=3-4 mice.

### Statistical Analysis

Statistical tests for behavior were calculated using GraphPad Prism version 10.6.0. Food intake, body weight, and activity were tested for significance using an unpaired t Test. Multiple comparisons were tested using an ordinary one-way ANOVA with Tukey’s multiple comparisons test. P values on all figures. Graphs were prepared using GraphPad Prism and figure layouts created using Adobe Illustrator.

---

**Supplemental Figure 1. Negative controls for dual-recombinase reporter activation.** (A–C) Representative images from *Ntsr1^Cre−^;Ntsr1^Flp+^*mice showing TH immunostaining (magenta) (A), mCherry expression (green) (B), and merged TH^+^ mCherry signal (C) in the ventral midbrain. (D–F) Additional caudal representative ventral midbrain images from *Ntsr1^Cre-^;Ntsr1^Flp+^*mice showing TH immunostaining (D), mCherry expression (E), and merged TH^+^ mCherry signal (F). (G–I) Representative images from *Ntsr1^Cre−^;Dat^Flp+^*mice showing TH immunostaining (G), mCherry expression (H), and merged TH^+^ mCherry signal (I) in the ventral midbrain. (J–L) Representative of additional caudal ventral midbrain images from *Ntsr1^Cre^;Dat^Flp^* mice showing TH immunostaining (J), mCherry expression (K), and merged TH^+^ mCherry signal (L). (M–O) Representative forebrain images from *Ntsr1^Cre^;Ntsr1^Flp^* mice showing DAPI labeling (blue) (M), mCherry expression (green) (N), and merged DAPI/mCherry signal (O). (P–R) Representative forebrain images from *Ntsr1^Cre^;Dat^Flp^* mice showing DAPI labeling (blue) (P), mCherry expression (green) (Q), and merged DAPI/mCherry signal (R). (S–U) Representative forebrain images from *Dat^Cre^;Dat^Flp^* mice showing DAPI labeling (blue) (S), mCherry expression (green) (T), and merged DAPI/mCherry signal (U). (V–X) Representative forebrain images from *Dat-^Cre^;Ntsr1^Flp^*mice showing DAPI labeling (blue) (V), mCherry expression (green) (W), and merged DAPI/mCherry signal (X). No recombinase-independent reporter expression was observed.

**Supplemental Figure 2. DAT and TH co-labeling in intersectionally targeted neurons.** Representative coronal sections from *Dat^Cre^*;*Dat^Flp^*, *Dat^Cre^*;*Ntsr1^Flp^*, and *Ntsr1^Cre^*;*Dat^Flp^*mice following systemic Con/Fon reporter delivery. TH (magenta), DAT (blue), and mCherry (green) are shown. (A,B) Representative images from *Dat^Cre^*;*Dat^Flp^*mice showing TH immunostaining (magenta), mCherry expression (green), and merged TH^+^ mCherry images in the substantia nigra (SN) (A) and ventral tegmental area (VTA) (B). Yellow arrowheads indicate TH⁺/mCherry⁺ cells. White arrowheads indicate mCherry⁺ cells. Open arrowheads indicate TH⁺ cells. (C,D) Representative images from *Dat^Cre^;Dat^Flp^* mice showing DAT immunostaining (blue), mCherry expression (green), and merged DAT^+^ mCherry images in the SN (C) and VTA (D) displayed in cyan. Yellow arrowheads indicate DAT⁺/mCherry⁺ cells. Red arrowheads indicate DAT⁺/mCherry⁻ cells. White arrowheads indicate mCherry⁺/DAT⁻ cells. (E,F) Representative images from *Dat^Cre^*;*Ntsr1^Flp^*mice showing TH immunostaining (magenta), mCherry expression (green), and merged TH^+^ mCherry images in the SN (E) and VTA (F). Yellow arrowheads indicate TH⁺/mCherry⁺ cells. White arrowheads indicate mCherry⁺ cells. Open arrowheads indicate TH⁺ cells. (G,H) Representative images from *Dat^Cre^*;*Ntsr1^Flp^*mice showing DAT immunostaining (blue), mCherry expression (green), and merged DAT^+^ mCherry images in the SN (G) and VTA (H). Yellow arrowheads indicate DAT⁺/mCherry⁺ cells. Red arrowheads indicate DAT⁺/mCherry⁻ cells. White arrowheads indicate mCherry⁺/DAT⁻ cells. (I,J) Representative images from *Ntsr1^Cre^*;*Dat^Flp^*mice showing TH immunostaining (magenta), mCherry expression (green), and merged TH^+^ mCherry images in the SN (I) and VTA (J). Yellow arrowheads indicate TH⁺/mCherry⁺ cells. White arrowheads indicate mCherry⁺ cells. Open arrowheads indicate TH⁺ cells. (K,L) Representative images from *Ntsr1^Cre^*;*Dat^Flp^*mice showing DAT immunostaining (blue), mCherry expression (green), and merged DAT + mCherry images in the SN (K) and VTA (L). Yellow arrowheads indicate DAT⁺/mCherry⁺ cells. Red arrowheads indicate DAT⁺/mCherry⁻ cells. White arrowheads indicate mCherry⁺/DAT⁻ cells. Insets show higher-magnification views of boxed regions. A small subset of TH⁻/DAT⁺/mCherry⁺ neurons were observed across all genotypes.

## Declaration of generative AI use

During the preparation of this work the author(s) used ChatGPT to edit the manuscript. After using this tool/service, the authors reviewed and edited the content as needed and take full responsibility for the content of the published article.

## Acknowledgements

We are grateful to Michigan State University’s Transgenic and Genome Editing Facility for generating the *NtsR12A-FlpO* knock-in mice, and in particular Drs. Elena Demireva and Huirong Xie. pAAV-Ef1a-fDIO mCherry and pAAV-nEF-Con/Fon TVA-mCherry were gifts from Karl Deisseroth (Stanford University). We are grateful to Dr. Timothy Shay and staff of the Caltech CLOVER center for advice on viral production.

