## Supplementary figures and images for "A Knock-in Ntsr1-Flp Driver Enables Intersectional and Systemic Targeting of Heterogeneous Midbrain Dopamine Circuits"

### Supplemental Figure 1

#### Supplemental Figure 1

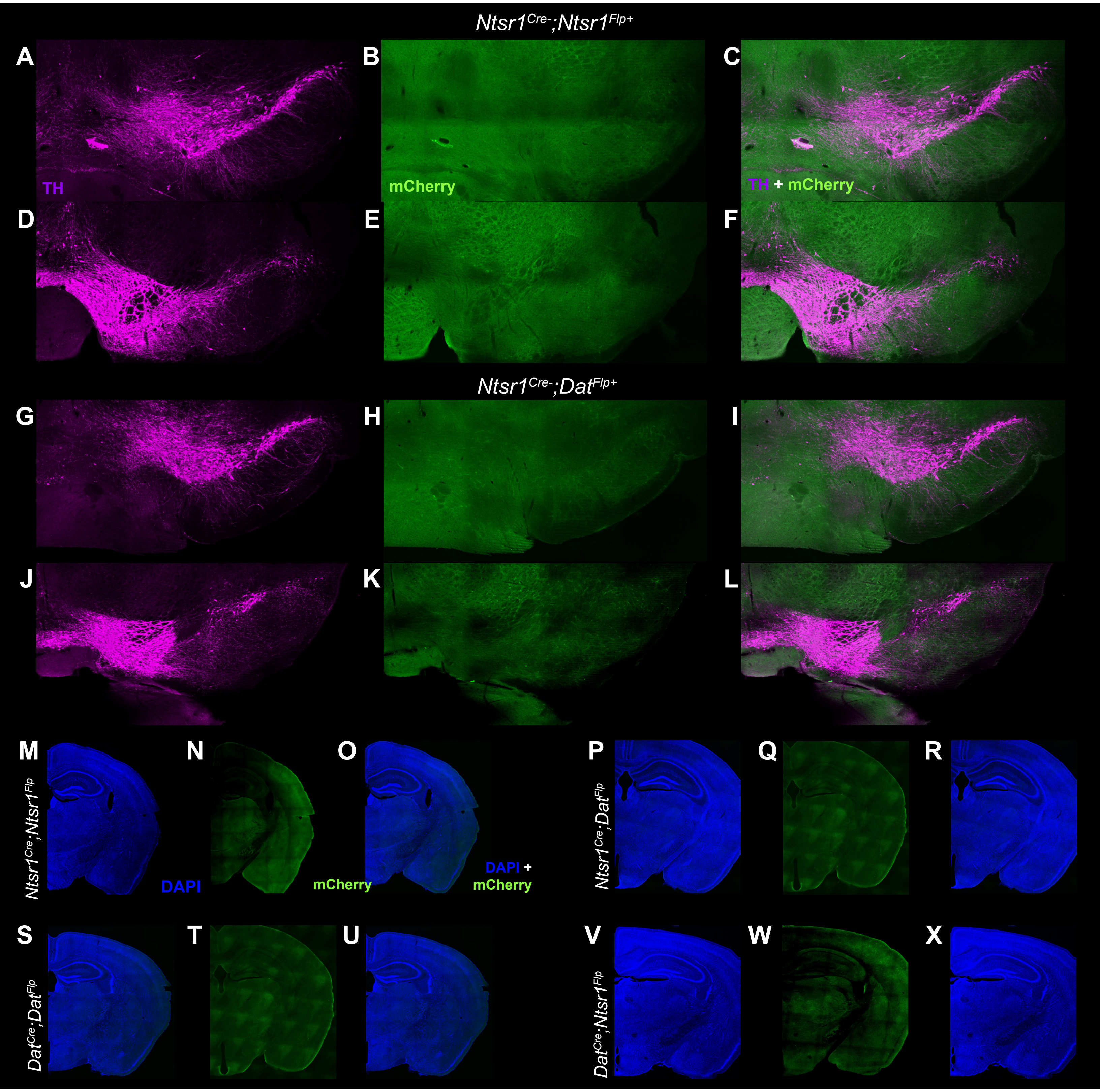

### Supplemental Figure 2

##### Supplemental Figure 2

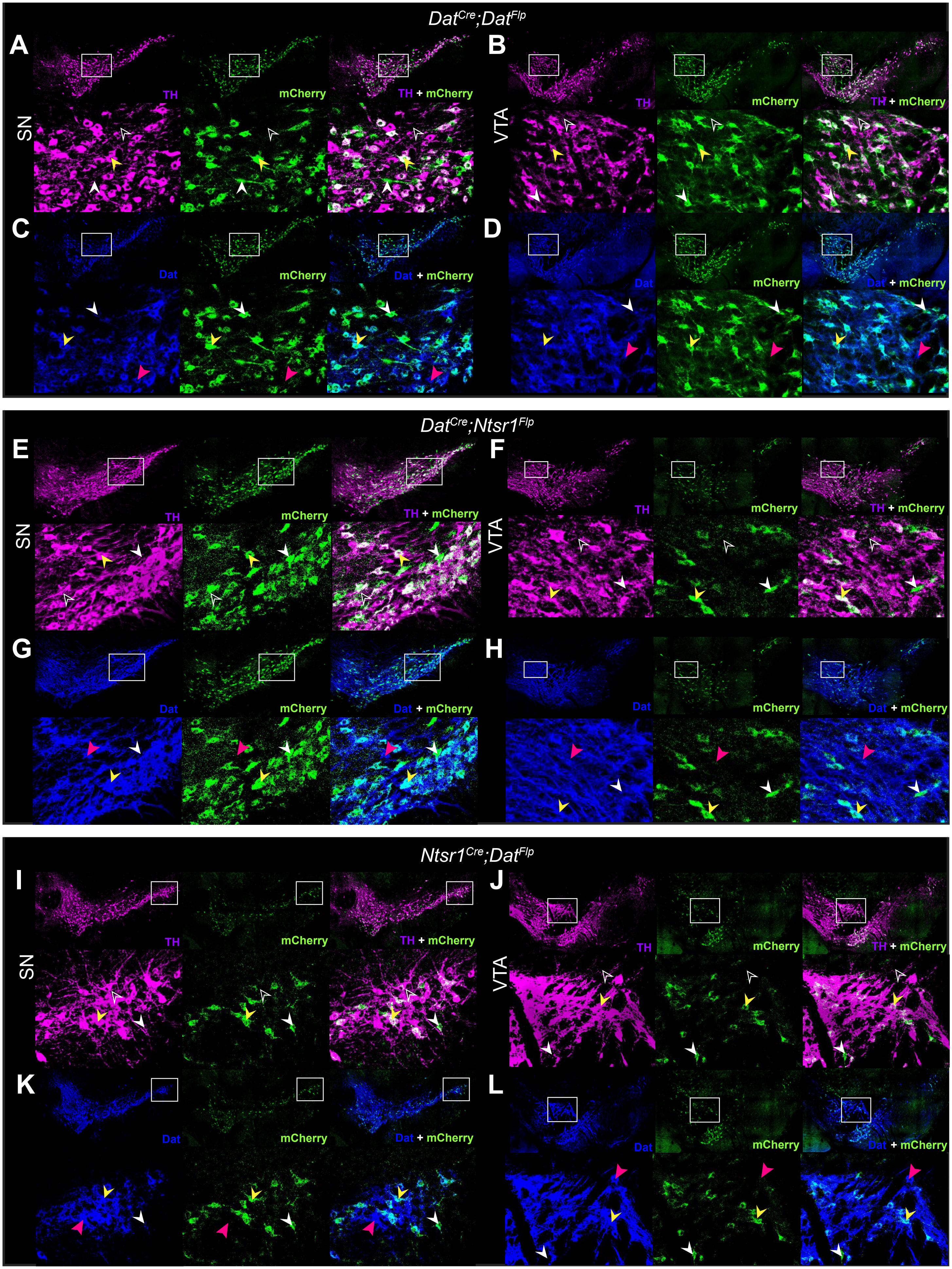
